# Trying to be unpredictable even against unobserved opponents shifts humans away from randomness but not from human predictability

**DOI:** 10.1101/2025.11.16.688665

**Authors:** Kagari Yamada, Kazushi Tsutsui, Kazutoshi Kudo

## Abstract

In competitive settings, randomization can prevent exploitation by opponents, yet human choices depend on both opponents’ past actions and players’ own action histories. Why this latter dependence arises, and how it affects predictability to others, remain unclear. Across three experiments, we separated sequence generation from subsequent prediction, thereby preventing generators from adapting to predictors’ responses. Whereas sequences generated under instructions to be random were biased toward alternation, those generated to evade prediction were biased toward repetition. Nevertheless, these sequences remained predictable. Human predictors systematically over-predicted repetitions, so sequences with a slight bias toward alternation actually minimized predictability. Thus, the failure to evade prediction may arise not merely from deviations from randomness, but also from inaccurate estimates of how others would predict one’s behavior. These findings suggest that understanding human competitive behavior requires considering not only how people adapt to others’ actions, but also the human-specific biases emerging in these processes.

## Introduction

Predicting what will happen next is a central function of human cognition^1–3^. In competitive social interactions, prediction is often bidirectional: opponents try to anticipate one another’s actions. Agents may therefore gain an advantage not only by anticipating others’ actions but also by making their own actions difficult to anticipate. Game theory provides a normative framework for addressing this challenge^4,5^. In two-player zero-sum games, a defensively optimal strategy is for players to randomize their choices so that no exploitable regularities remain. Such mixed-strategy equilibria can partially explain human behavior^6,7^. Nevertheless, human choices in repeated competitive settings are influenced by both the opponent’s past actions^8,9^ and their own action histories^10–14^.

Extensive research has characterized how people learn statistical regularities in external event sequences^15–18^ and has described how players use opponents’ past actions to adjust their choices in pursuit of higher payoffs^19,20^. By contrast, even though randomization is the normative solution, less is known about what players’ dependence on their own action histories reflects when they try to evade prediction. One possibility is that people intend to implement objective randomization but rely on a distorted representation of randomness. When people attempt to behave randomly, they systematically deviate from objective randomness^21–24^. Specifically, they tend to over-alternate, producing fewer repetitions than expected by chance^25,26^. Thus, departures from mixed-strategy play in competitive environments may partly reflect a failure to implement the very randomization needed to render behavior objectively unpredictable.

Another possibility is that people do not aim for objective randomness at all. Equilibrium randomization shields a player from exploitation but need not maximize gains against a systematically biased opponent; exploiting such a bias may require deviating from equilibrium play^27,28^. This distinction is important because human observers are sensitive to sequence structures at multiple levels, including marginal item frequencies and transition probabilities^16,17,29,30^; however, their predictions are not perfectly calibrated to objective sequence statistics^31–33^. Notably, such biases could provide a target for prediction evasion, even when generators cannot observe a specific predictor’s responses. Rather than adapting to an individual, people may generate non-random sequences based on general expectations about how others predict.

In this study, we temporally dissociated sequence generation from subsequent prediction, ensuring that generators had no access to the trial-by-trial responses of the participants who later predicted their sequences. This design allowed us to examine dependence on generators’ own action histories without feedback-driven adaptation to a specific predictor and to assess how predictable the resulting sequences were to independent observers. Participants generated sequences with a distinctive non-random structure. Rather than displaying the over-alternation typically observed under instructions to be random, prediction-evasion sequences were biased toward repetition. Nevertheless, this generation pattern failed to reduce predictability. Human observers tracked transition statistics but systematically predicted repetitions more often than they occurred. The generators’ repetition bias therefore aligned with, rather than counteracted, the observers’ tendency to predict repetition. The results indicate that attempts at strategic unpredictability failed not merely because the generators departed from objective randomness but also because their generated sequences were not appropriately calibrated to how human observers predicted them. By separating objective randomness from human predictability, this study links sequence cognition with strategic interaction and suggests that attempts to outguess others can fail when behavior is guided by miscalibrated assumptions about how others predict.

## Results

### Intention to evade prediction induces a repetition bias

We first asked whether the goal of evading another person’s prediction leads people to approximate objective randomization, or instead produces a distinct sequence structure. We examined sequence generation in the absence of any information about the opponent to isolate this goal from feedback-based adaptation. Specifically, we compared sequences generated to approximate randomness with those generated to avoid another person’s trial-by-trial predictions.

In Experiment 1, the participants generated a sequence of binary choices (left or right) over 10 blocks of 20 trials (Fig. 1A, B). They were randomly assigned to receive one of two instructions. In the competitive instruction, participants (n = 25) were asked to generate a sequence that would be difficult for later participants to predict on a trial-by-trial basis. In the random instruction, participants (n = 25) were asked to generate a sequence that was as random as possible. We refer to the resulting sequence classes as Competitive and Random sequences, respectively; these labels denote the generation instruction and not the actual predictability or randomness of the sequences.

**Figure 1.**
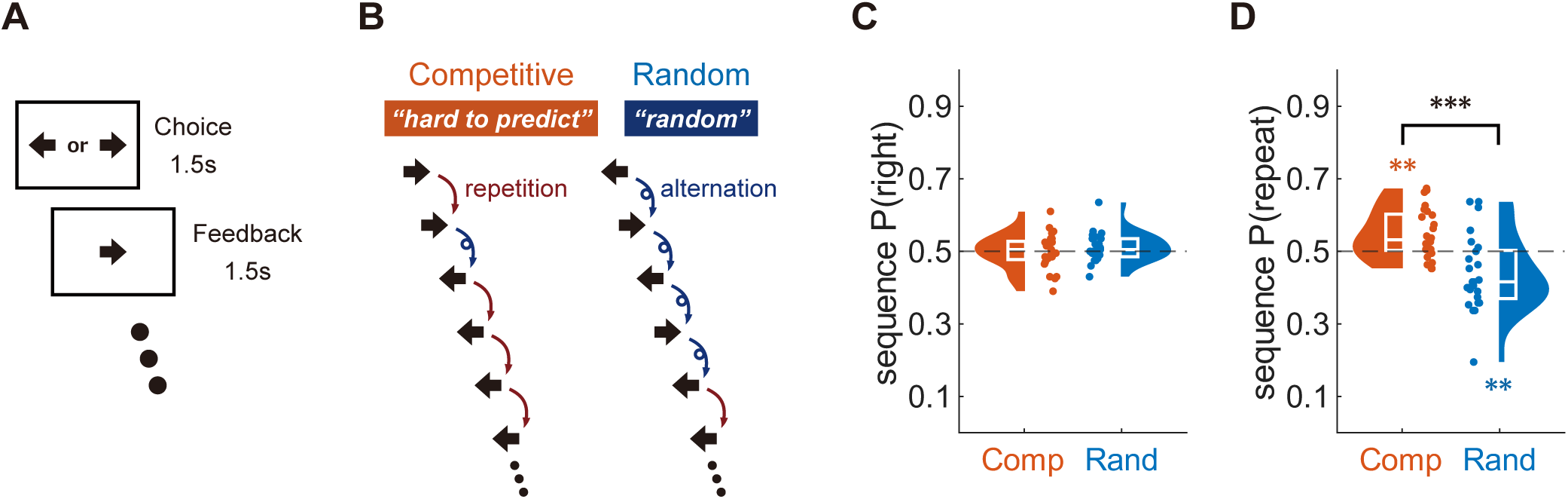
Intention to evade prediction induces a repetition bias. (A) Example trial structure in Experiment 1. Participants generated binary choices over 10 blocks of 20 trials. Each trial comprised a choice phase followed by feedback indicating the selected item. After each block, participants rated how well they kept the task instruction in mind. (B) Schematic examples of the two generation instructions and resulting transition structures. In the competitive instruction, participants generated sequences intended to be difficult for a later participant to predict on a trial-by-trial basis. In the random instruction, participants generated sequences intended to be as random as possible. These sequence classes are referred to as Competitive and Random sequences, respectively. A repetition occurs when a choice matches the immediately preceding choice; an alternation occurs when it differs from the immediately preceding choice. (C) Marginal side frequency in generated sequences. Half-violin plots reveal participant-level sequence P(right), defined as the probability that an item in the generated sequence was rightward, for Competitive and Random sequences. Embedded box plots present the median and interquartile range; dots indicate individual participants. The horizontal dashed line indicates balanced side use, sequence P(right) = 0.5. (D) First-order transition structure of generated sequences. Half-violin plots reveal participant-level sequence P(repeat), defined as the probability that a sequence item repeated the immediately preceding item, for Competitive and Random sequences. The horizontal dashed line signifies chance-level transition balance, sequence P(repeat) = 0.5. Black asterisks denote between-sequence class differences and red and blue asterisks denote one-sample differences from chance within competitive and random sequences, respectively. \*\**p* < 0.01, \*\*\**p* < 0.001.

First, we asked whether the two sequence classes differed in their marginal use of left and right choices. Here, the sequence P(right) denotes the probability that an item in the generated sequence was rightward. The overall sequence P(right) did not differ reliably between Competitive and Random sequences (*t* (48) = -0.87, *p* = 0.387, Cohen’s *d* (*d*) = - 0.25, 95% confidence interval (CI) [-0.82, 0.32]) (Fig. 1C). The absolute magnitude of side bias, |sequence P(right) − 0.5|, was also comparable between Competitive and Random sequences (*U* = 359, *p* = 0.370, *r* = 0.13, CI [-0.16, 0.39]) (Fig. S1C). This absence of differential side preference was stable across the task: block-wise analysis revealed no reliable main effect of block (*F* (6.75, 324.18) = 1.40, *p* = 0.205, *η_G_²* = 0.024) or instruction × block interaction (*F* (6.75, 324.18) = 0.73, *p* = 0.641, *η_G_²* = 0.013) (Fig. S1A).

However, the two sequence classes diverged in their first-order transition structure (Fig. 1D). We quantified this structure as sequence P(repeat), which is the probability that a sequence item repeated the immediately preceding item. Random sequences showed a typical alternation bias^25,26^: their sequence P(repeat) was significantly below the 0.5 chance level (*t* (24) = -3.01, *p* = 0.006, *d* = -0.60, CI [-1.05, -0.15]). Competitive sequences showed the opposite pattern: their sequence P(repeat) was significantly > 0.5, indicating a repetition bias (*t* (24) = 3.52, *p* = 0.002, *d* = 0.70, CI [0.24, 1.17]). A direct comparison confirmed that the repetition probability was substantially higher for Competitive than for Random sequences (*t* (48) = 4.44, *p* < 0.001, *d* = 1.26, CI [0.63, 1.88]). This transition-level difference was stable across blocks: block-wise analysis revealed a robust main effect of instruction (*F* (1, 48) = 19.71, *p* < 0.001, *η_G_²* = 0.126), with no reliable main effect of block (*F* (6.51, 312.45) = 0.75, *p* = 0.618, *η_G_²* = 0.010) or instruction × block interaction (*F* (6.51, 312.45) = 1.20, *p* = 0.305, *η_G_²* = 0.016) (Fig. S1B).

When deviation from objective first-order randomness was assessed irrespective of direction, Competitive sequences were closer to transition balance than Random sequences (*U* = 198, *p* = 0.027, *r* = -0.31, CI [-0.54, -0.04]) (Fig. S1D); however, this deviation did not disappear but instead formed a repetition bias. These structural differences were unlikely to reflect differences in general task engagement, as mean self-reported engagement did not differ reliably between the two instruction conditions (*t* (48) = -1.22, *p* = 0.230, *d* = -0.34, CI [-0.92, 0.23]) (Fig. S2A).

Together, these results indicate that the goal of evading another person’s prediction changes sequence generation, even without observing the predictor’s responses. While Random sequences showed a canonical over-alternation bias, Competitive sequences crossed the transition balance and became repetition-biased. Thus, the goal of strategic unpredictability did not produce objective randomization, but a distinct non-random generation pattern centered on repetition.

### Sequences intended to evade prediction remain vulnerable to human observers

In Experiment 2, we directly tested whether the sequences generated with the objective of difficulty in prediction were resistant to human prediction. All the 50 sequences generated in Experiment 1 were used as target sequences: 25 Competitive sequences and 25 Random sequences. A new group of participants (n = 250), none of whom were involved in Experiment 1, predicted these sequences trial by trial, with five participants assigned to each sequence (Fig. 2A, B). To avoid biasing their predictions^31^, the participants were provided no information about the origin or generative process of the sequences they were asked to predict.

**Figure 2.**
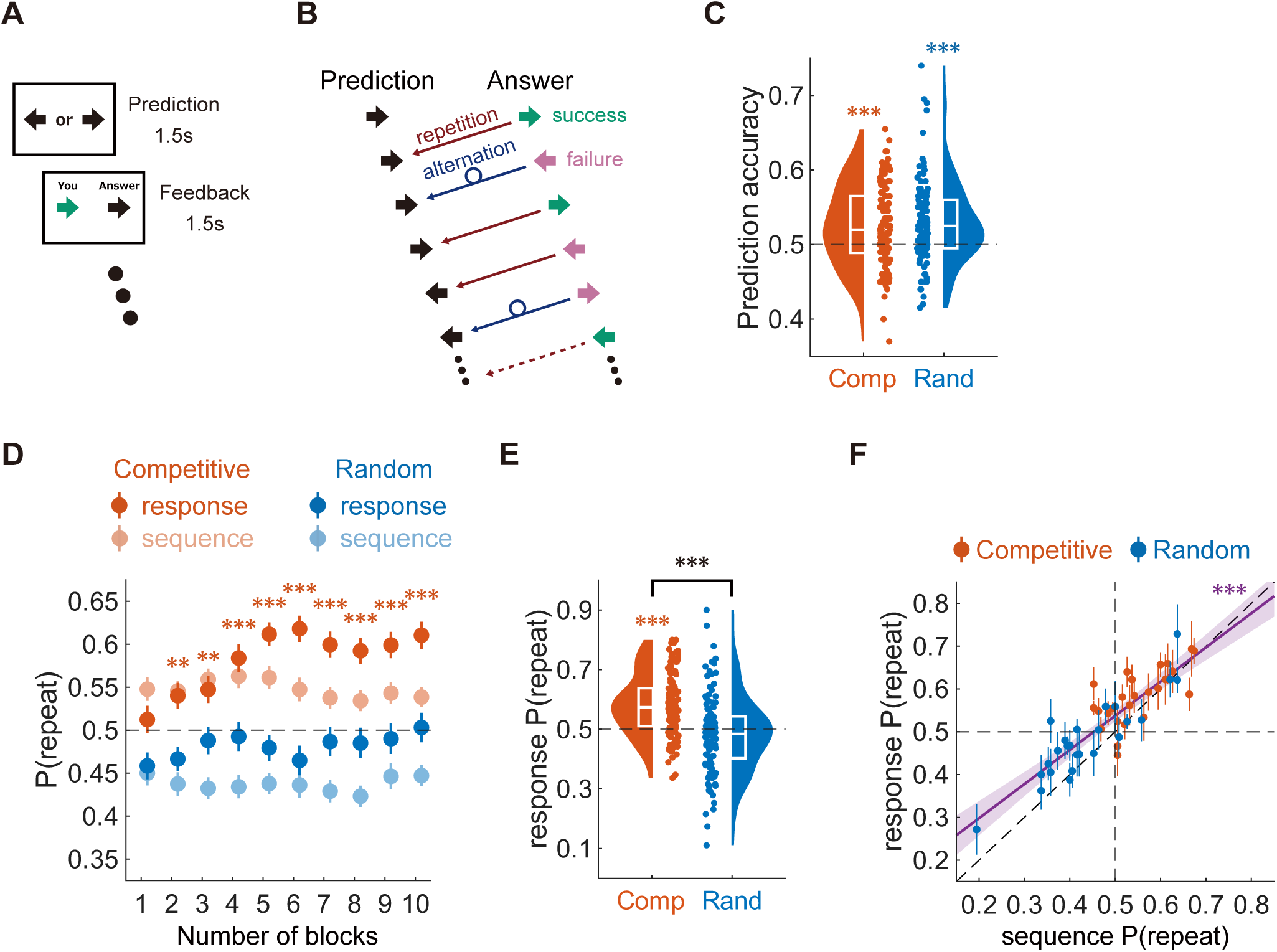
Human-generated sequences intended to evade prediction remain predictable. (A) Example trial structure in Experiment 2. Participants predicted the next item in a binary target sequence over 10 blocks of 20 trials. Each trial comprised a prediction phase followed by feedback showing the participant’s prediction and accurate target item. (B) Definition of prediction strategies. A repetition prediction signifies that the participant predicted the next item to match the immediately preceding target item; an alternation prediction shows that the participant predicted the opposite item. Prediction accuracy was defined by whether the prediction matched the current target item. (C) Prediction accuracy for Competitive and Random sequences. Half-violin plots illustrate participant-level accuracy; embedded box plots show the median and interquartile range; dots indicate individual participants. The horizontal dashed line indicates the chance level. (D) Block-wise sequence and response repetition probabilities. Sequence P(repeat) denotes the probability that the target item repeated the immediately preceding target item. Response P(repeat) signifies the probability that participants predicted such a repetition. Light-colored points display sequence P(repeat); dark-colored points demonstrate response P(repeat). Error bars indicate ±1 SEM. The horizontal dashed line indicates the chance-level repetition probability, P(repeat) = 0.5. (E) Overall response P(repeat), collapsed across blocks. Values > 0.5 indicate repetition expectation; values < 0.5 indicate alternation expectation. (F) Relationship between sequence P(repeat) and response P(repeat). Each point represents one target sequence; error bars indicate ±1 SEM across the five participants assigned to the particular sequence. The purple line presents the linear fit across all sequences; shading indicates the 95% confidence interval. The diagonal dashed line denotes the identity line; horizontal and vertical dashed lines indicate P(repeat) = 0.5. Black asterisks denote between-sequence class differences and red and blue asterisks signify one-sample differences from chance within Competitive and Random sequences, respectively. P-values were corrected using the false discovery rate procedure for block-wise analyses. \*\**p* < 0.01, \*\*\**p* < 0.001.

Contrary to the generators’ goal, Competitive sequences were predicted significantly above the 50% chance level (*t* (124) = 5.12, *p* < 0.001, *d* = 0.46, CI [0.27, 0.64]) (Fig. 2C). Random sequences were also predicted above chance (*t* (124) = 5.93, *p* < 0.001, *d* = 0.53, CI [0.34, 0.72]), and there was no evidence that Competitive sequences were more difficult to predict than Random sequences (*t* (248) = -0.59, *p* = 0.556, *d* = -0.07, CI [-0.32, 0.17]). Thus, sequences generated with the explicit goal of making them difficult for another person were nevertheless predicted above chance by independent human observers.

Next, we examined how participants achieved the above-chance prediction. As Experiment 1 revealed that Competitive sequences were biased toward repetition whereas Random sequences were biased toward alternation, we examined whether the participants’ predictions reflected these transition statistics. Here, sequence P(repeat) denotes the probability that the target item repeated the immediately preceding target item, whereas response P(repeat) denotes the probability that the participants predicted such a repetition.

At the block level, the prediction patterns initially suggested asymmetric learning of the two sequence classes. For Competitive sequences, the response P(repeat) increased above chance and broadly followed a repetition-favoring sequence structure. However, for Random sequences, the response P(repeat) remained close to chance despite the alternation-favoring structure of the sequences (Fig. 2D). This pattern was preserved when predictions were collapsed across blocks: participants predicting Competitive sequences showed a robust repetition expectation (*t* (124) = 8.94, *p* < 0.001, *d* = 0.80, CI [0.60, 1.00]), whereas participants predicting Random sequences did not demonstrate a reliable alternation expectation (*t* (124) = -1.59, *p* = 0.115, *d* = -0.14, CI [-0.32, 0.04]) (Fig. 2E).

Nonetheless, this apparent asymmetry should not be interpreted as a simple one-to-one mapping in which repetition-favoring Competitive sequences were learned whereas alternation-favoring Random sequences were not. A more parsimonious account is that participants tracked sequence P(repeat) but overestimated the probability that the same item would be repeated. Consistently, response P(repeat) exceeded sequence P(repeat) for both Competitive (*t* (124) = 3.94, *p* < 0.001, *d* = 0.35, CI [0.17, 0.53]) and Random sequences (*t* (124) = 4.80, *p* < 0.001, *d* = 0.43, CI [0.24, 0.61]) (Fig. S3A). Across the 50 target sequences, sequence P(repeat) strongly predicted response P(repeat) (*r* = 0.874, *p* < 0.001) (Fig. 2F), indicating that the participants were sensitive to transition statistics across both sequence classes. However, the fitted regression line shifted upward: at a sequence P(repeat) of 0.5, the fitted response P(repeat) was significantly greater than chance (estimated P(repeat) = 0.537, CI [0.524, 0.551]; *t* (48) = 5.70, *p* < 0.001, *d* = 0.81, CI [0.48, 1.14]). Thus, the weak expression of alternation in Random sequences did not reflect a categorical failure to learn those sequences; rather, it reflected a general tendency to over-predict repetition relative to the true transition probability.

This transition-based account was not the only possible source of predictability in the original human-generated sequences. The participants’ rightward predictions also tracked the marginal probability of the rightward target items (*r* = 0.530, *p* < 0.001) (Fig. S3B), and the marginal rightward probability and repetition probability were not easily separable across the original sequence set (Competitive: *r* = -0.389, *p* = 0.054; Random: *r* = 0.345, *p* = 0.091) (Fig. S3C). Therefore, Experiment 2 revealed that Competitive sequences were predicted above chance and that participants’ predictions tracked the transition statistics of their assigned sequences. However, as the original sequences also contained variations in the marginal side frequencies, Experiment 2 alone could not confirm whether the prediction was driven by transition statistics, marginal side frequencies, or both.

### Over-prediction of repetition persists even when side frequencies are balanced

Experiment 2 revealed that participants tracked sequence P(repeat), but also systematically over-predicted repetition relative to the true transition probability of the assigned sequence. Nevertheless, the original human-generated sequences also varied in their marginal left/right frequencies, leaving open the question of whether this repetition-biased calibration reflected transition statistics, side-frequency information, or both. We generated a new set of synthetic sequences for Experiment 3 to isolate the contribution of the transition structure. For each original sequence, we preserved its first-order transition structure and closely matched its higher-order run length statistics, while balancing the marginal side frequencies (Fig. 3A). We refer to synthetic sequences derived from the original Competitive and Random sequences as Competitive-like and Random-like sequences, respectively; these labels indicate the source transition structure, whereas all synthetic sequences were balanced in marginal side frequency. Subsequently, a new group of participants (n = 250) predicted these balanced synthetic sequences trial by trial, with five participants assigned to each sequence.

**Figure 3.**
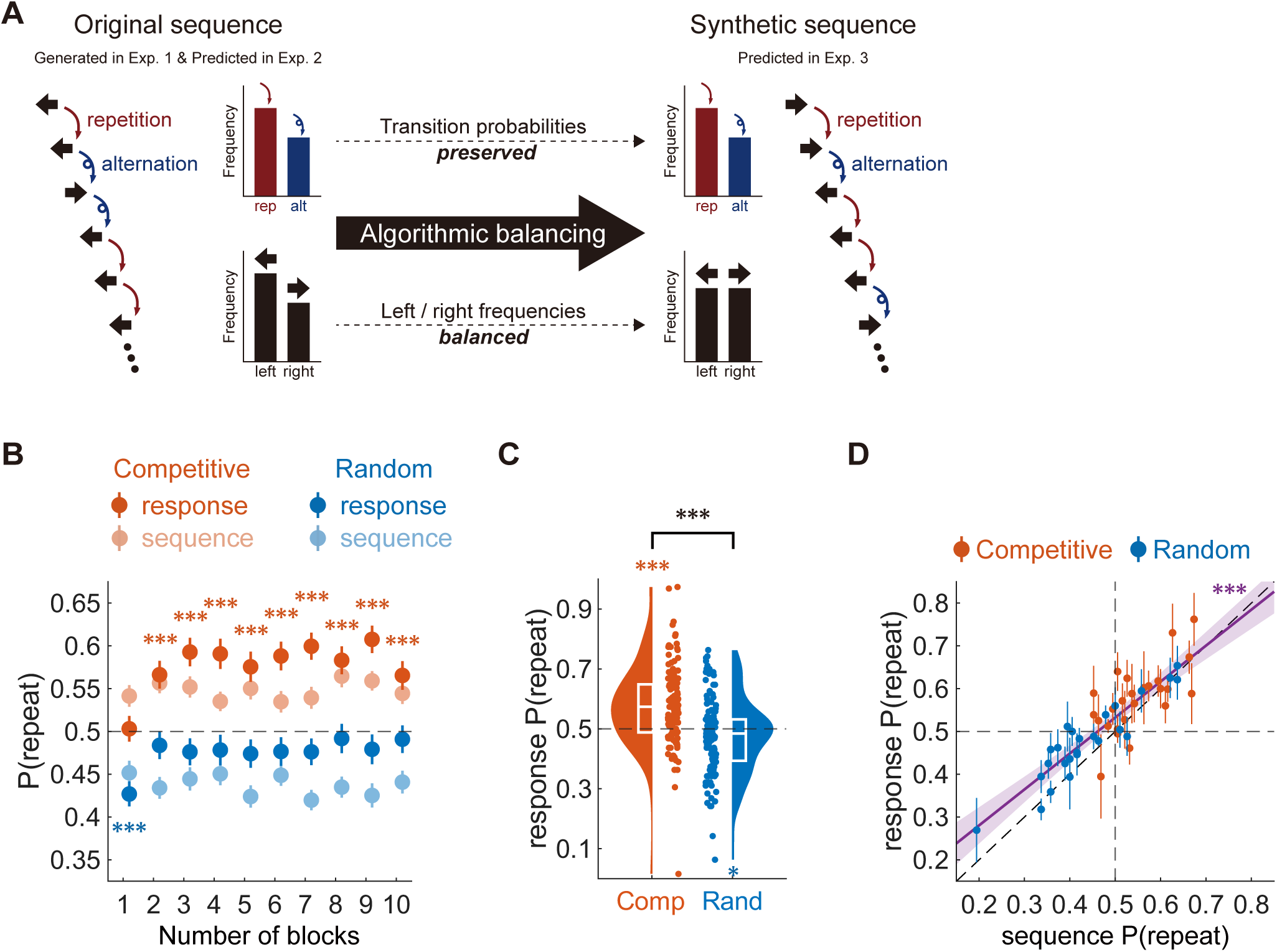
Balanced marginal frequencies revealing persistent over-prediction of repetition. (A) Schematic of stimulus construction for Experiment 3. Original human-generated sequences from Experiment 1, predicted in Experiment 2, were transformed into synthetic sequences. The transformation preserved the transition structure of each sequence, including its repetition/alternation profile, while balancing the marginal frequencies of left and right items. The synthetic sequences were subsequently employed as target sequences in Experiment 3. (B) Block-wise sequence and response repetition probabilities in Experiment 3. Sequence P(repeat) indicates the probability that the target item repeated the immediately preceding target item. Response P(repeat) denotes the probability that participants predicted such a repetition. Light-colored points display sequence P(repeat); dark-colored points present response P(repeat). Error bars indicate ±1 SEM. The horizontal dashed line signifies the chance-level repetition probability, P(repeat) = 0.5. (C) Overall response P(repeat), collapsed across blocks. Half-violin plots display participant-level response P(repeat); embedded box plots display the median and interquartile range; dots indicate individual participants. Values > 0.5 indicate repetition expectation; values < 0.5 indicate alternation expectation. (D) Relationship between sequence P(repeat) and response P(repeat) in Experiment 3. Each point represents one balanced target sequence; error bars indicate ±1 SEM across the five participants assigned to the particular sequence. The purple line presents the linear fit across all sequences; shading indicates the 95% confidence interval. The diagonal dashed line denotes the identity line; horizontal and vertical dashed lines signify P(repeat) = 0.5. Black asterisks denote between-sequence class differences and red and blue asterisks signify one-sample differences from chance within Competitive-like and Random-like sequences, respectively. P-values were corrected using the false discovery rate procedure for block-wise analyses. \**p* < 0.05, \*\*\**p* < 0.001.

The balanced sequences reproduced the key transition-based prediction pattern observed in Experiment 2. At the block level, response P(repeat) tracked the broad separation between Competitive-like and Random-like sequences, despite the absence of marginal side-frequency cues (Fig. 3B). When collapsed across blocks, participants predicting Competitive-like sequences showed a robust repetition expectation (*t* (124) = 6.55, *p* < 0.001, *d* = 0.59, CI [0.39, 0.78]), whereas participants predicting Random-like sequences demonstrated only a weaker alternation expectation (*t* (124) = - 2.18, *p* = 0.032, *d* = -0.19, CI [-0.37, -0.02]) (Fig. 3C). Thus, the balancing left and right frequencies did not eliminate the asymmetric expression of repetition-versus alternation-favoring structures.

However, as in Experiment 2, this apparent asymmetry was better explained as biased calibration than as a categorical failure to learn Random-like sequences. Response P(repeat) exceeded sequence P(repeat) in both Competitive-like (*t* (124) = 2.69, *p* = 0.008, *d* = 0.24, CI [0.06, 0.42]) and Random-like sequences (*t* (124) = 4.41, *p* < 0.001, *d* = 0.39, CI [0.21, 0.58]) (Fig. S4A). Across the 50 balanced sequences, sequence P(repeat) strongly predicted response P(repeat) (*r* = 0.873, *p* < 0.001) (Fig. 3D). Notably, the fitted regression line again shifted upward, and at a sequence P(repeat) of 0.5, the fitted response P(repeat) was significantly greater than chance (estimated P(repeat) = 0.533, CI [0.519, 0.547]; *t* (48) = 4.68, *p* < 0.001, *d* = 0.66, CI [0.35, 0.98]). These results indicate that participants remained sensitive to transition statistics when side frequencies were uninformative but continued to overestimate the probability that the same item was repeated.

Finally, we compared sequence-level prediction patterns between the original sequences in Experiment 2 and their balanced counterparts in Experiment 3. The response P(repeat) was strongly aligned between the two experiments (*r* = 0.795, *p* < 0.001) (Fig. S4B), indicating that the transition-based component of the prediction was preserved after removing the marginal side frequencies. By contrast, the response P(right) no longer showed a reliable relationship between the two experiments (*r* = 0.008, *p* = 0.957) (Fig. S4C). Hence, marginal side frequencies influenced side-specific predictions in Experiment 2, but the repetition-prediction pattern reflected sensitivity to the transition structure. Overall, Experiments 2 and 3 reveal that human sequence prediction is shaped by both marginal occurrence and transition statistics; however, the over-prediction of repetition persists even when the marginal side frequency is no longer informative.

### Biased calibration toward repetition generalizes across higher-order sequence contexts

The preceding analyses confirmed that the participants tracked first-order transition statistics but systematically overestimated the probability of repetition. However, humans can also learn higher-order probabilities in sequential contexts^34,35^. Further, classic studies suggest that human sequence prediction biases can invert depending on the run length—for example, expecting alternation after short runs but repetition after long runs^36^. Hence, the overestimation of repetition we observed might be particular to certain run lengths, or participants might exhibit entirely different response tendencies in higher-order contexts. To investigate this possibility, we examined run-length-conditioned statistics: the probability of a target repeating after 1–4 consecutive identical items (Fig. 4A).

**Figure 4.**
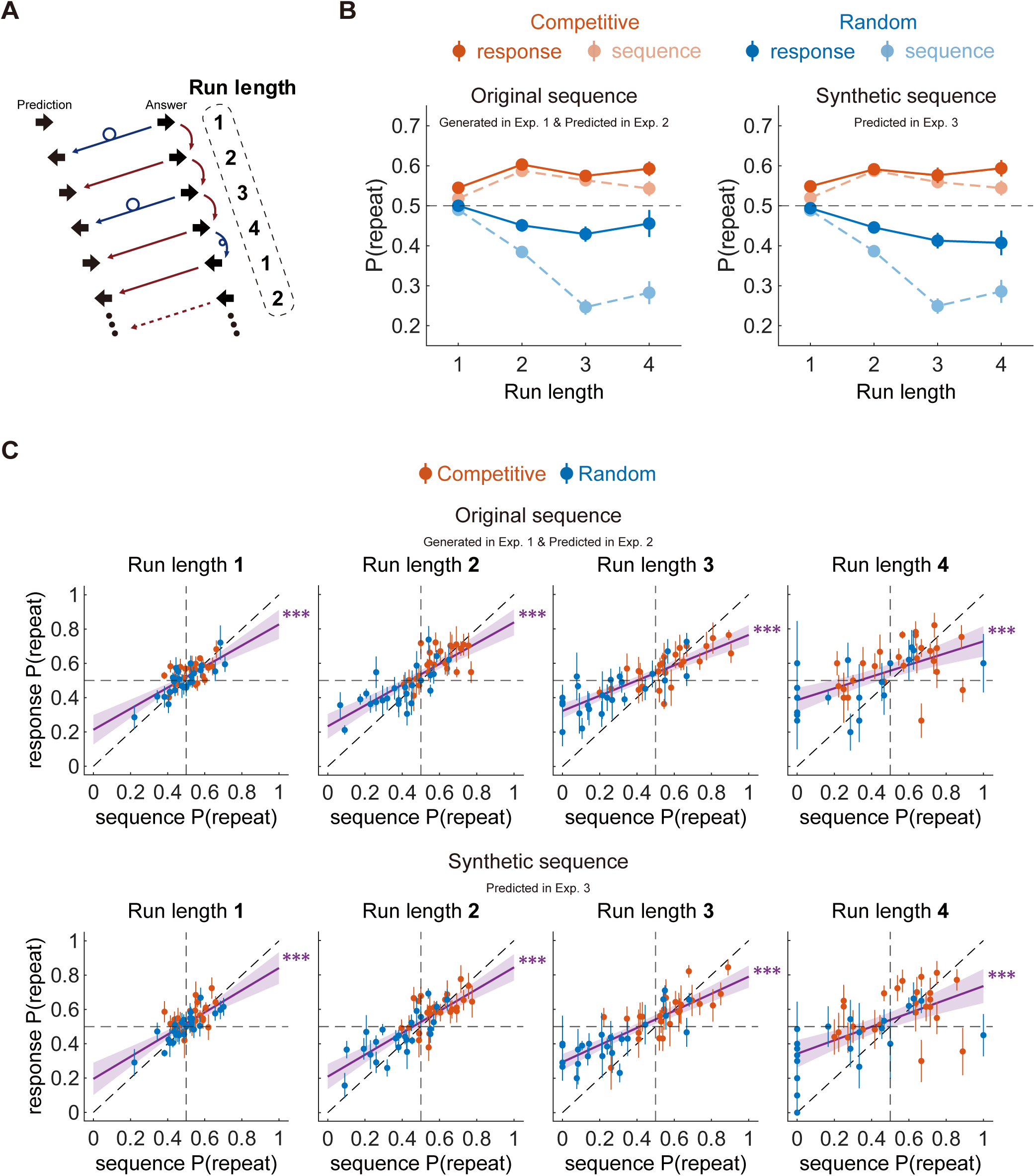
Higher-order sequence tracking systematically calibrated toward repetition. (A) Definition of run length in the prediction task. Run length signifies the number of consecutive identical target items immediately preceding the current prediction. A repetition prediction indicates that the participant predicted the next item to match the immediately preceding target item; an alternation prediction denotes that the participant predicted the opposite item. (B) Run length-conditioned sequence and response repetition probabilities. The left panel presents original human-generated sequences from Experiment 1, predicted in Experiment 2; the right panel shows the corresponding balanced synthetic sequences predicted in Experiment 3. Sequence P(repeat) denotes the probability that the target item repeated after a given run length. Response P(repeat) captures the probability that participants predicted such a repetition. Light-colored dashed lines show sequence P(repeat); dark-colored solid lines display response P(repeat). Error bars indicate ±1 SEM. The horizontal dashed line denotes the chance-level repetition probability, P(repeat) = 0.5. (C) Run length-conditioned sequence P(repeat) predicts response P(repeat). Each subpanel reveals one run length. The upper row shows original sequences predicted in Experiment 2; the lower row depicts balanced synthetic sequences predicted in Experiment 3. Each point represents one target sequence; error bars indicate ±1 SEM across the five participants assigned to the particular sequence. The purple line denotes the linear fit across all sequences; shading indicates the 95% confidence interval. The diagonal dashed line presents the identity line; horizontal and vertical dashed lines indicate P(repeat) = 0.5. Red and blue indicate Competitive/Competitive-like and Random/Random-like sequences, respectively. Purple asterisks denote the significance of the overall sequence-level relationship between sequence P(repeat) and response P(repeat). \*\*\**p* < 0.001.

First, we investigated whether participants’ run length-conditioned predictions tracked the actual higher-order structures of the assigned sequences. In both the original (Experiment 2) and balanced synthetic sequences (Experiment 3), which preserved the run length-conditioned transition statistics of the original sequences while balancing the marginal left/right frequencies (Fig. S5A), the response P(repeat) broadly followed the empirical sequence P(repeat) across all run lengths (Fig. 4B). Thus, the participants demonstrated clear sensitivity to local higher-order structures across varying contexts.

Although the participants successfully tracked these local higher-order structures, their predictions remained subject to a persistent calibration bias. We analyzed the relationship between sequence P(repeat) and response P(repeat) for each run length (Fig. 4C) to test whether the consistent overestimation of repetition was generalizable to higher-order contexts. Across run lengths 1–4, the run length-specific repetition probability of the sequence strongly predicted the participants’ corresponding repetition predictions in both Experiments 2 (*r* = 0.604-0.821, *ps* < 0.001) and 3 (*r* = 0.614-0.824, *ps* < 0.001). Notably, the fitted regression lines systematically shifted upward across all run lengths. At a sequence P(repeat) of 0.5, the fitted response P(repeat) was significantly above the chance level for all run lengths in Experiment 2 (estimated P(repeat) = 0.520-0.557; *t* (43-48) = 2.61-3.71, *ps* ≤ 0.012, *d* = 0.37-0.56) and for run lengths 1–3 in Experiment 3 (estimated P(repeat) = 0.519-0.542; *t* (48) = 2.24-3.23, *ps* < 0.030, *d* = 0.32-0.49), with the same positive tendency at run length 4 (estimated P(repeat) = 0.539, CI [0.494, 0.583]; *t* (43) = 1.75, *p* = 0.087, *d* = 0.27, CI [-0.04, 0.58]). This pervasive overestimation was further confirmed by directly evaluating the sequence-referenced deviations in the predictions for each run length (Fig. S5B); participants showed no statistical evidence of reversing their bias into an underestimation of repetition at any specific run length.

Finally, the run length-conditioned response, P(repeat), was preserved between the original sequences in Experiment 2 and their balanced synthetic counterparts in Experiment 3 (Fig. S5C). As the synthetic sequences preserved the higher-order transition structure while removing marginal side frequency cues, this cross-experimental alignment indicates that predictions continued to track higher-order sequence histories even when the marginal side frequency was uninformative. Together, these analyses reveal that observers tracked the higher-order transition structure but remained systematically calibrated toward repetition.

### Blind spot for human prediction is shifted toward alternation

In the preceding sections, we explained that human prediction is not calibrated to objective transition probabilities. Participants tracked the sequence P(repeat), but their response P(repeat) consistently shifted upward, indicating a tendency to over-predict repetition. This raises a different question regarding the sequences generated in Experiment 1. We initially evaluated these sequences relative to objective randomness, defined by balanced side frequency and transition balance. Yet, if human prediction is systematically biased, the sequence statistics defining objective randomness need not coincide with those minimizing predictability to human observers.

If human prediction lacked an overall repetition bias, the prediction line relating sequence P(repeat) to response P(repeat) would pass through the neutral point (0.5, 0.5) without an upward shift (dashed purple line in Fig. 5A). Under such unshifted calibration, the direction of the transition-based prediction would correctly align with the sequence structure across the full range: repetition-biased sequences would elicit repetition predictions, and alternation-biased sequences would produce alternation predictions. Consequently, the objective transition balance (sequence P(repeat) = 0.5) would be a natural candidate for minimal predictability. However, as the actual response P(repeat) is systematically shifted upward (solid purple line in Fig. 5A), the prediction curve crosses into a mismatch region (pink area in Fig. 5A), where sequence P(repeat) is < 0.5 but the response P(repeat) remains > 0.5. In this mismatch zone, the sequence is biased toward alternation, whereas the observer expects repetition. Therefore, such sequences may be more difficult to predict than objectively random sequences and may even drive below-chance prediction accuracy.

**Figure 5.**
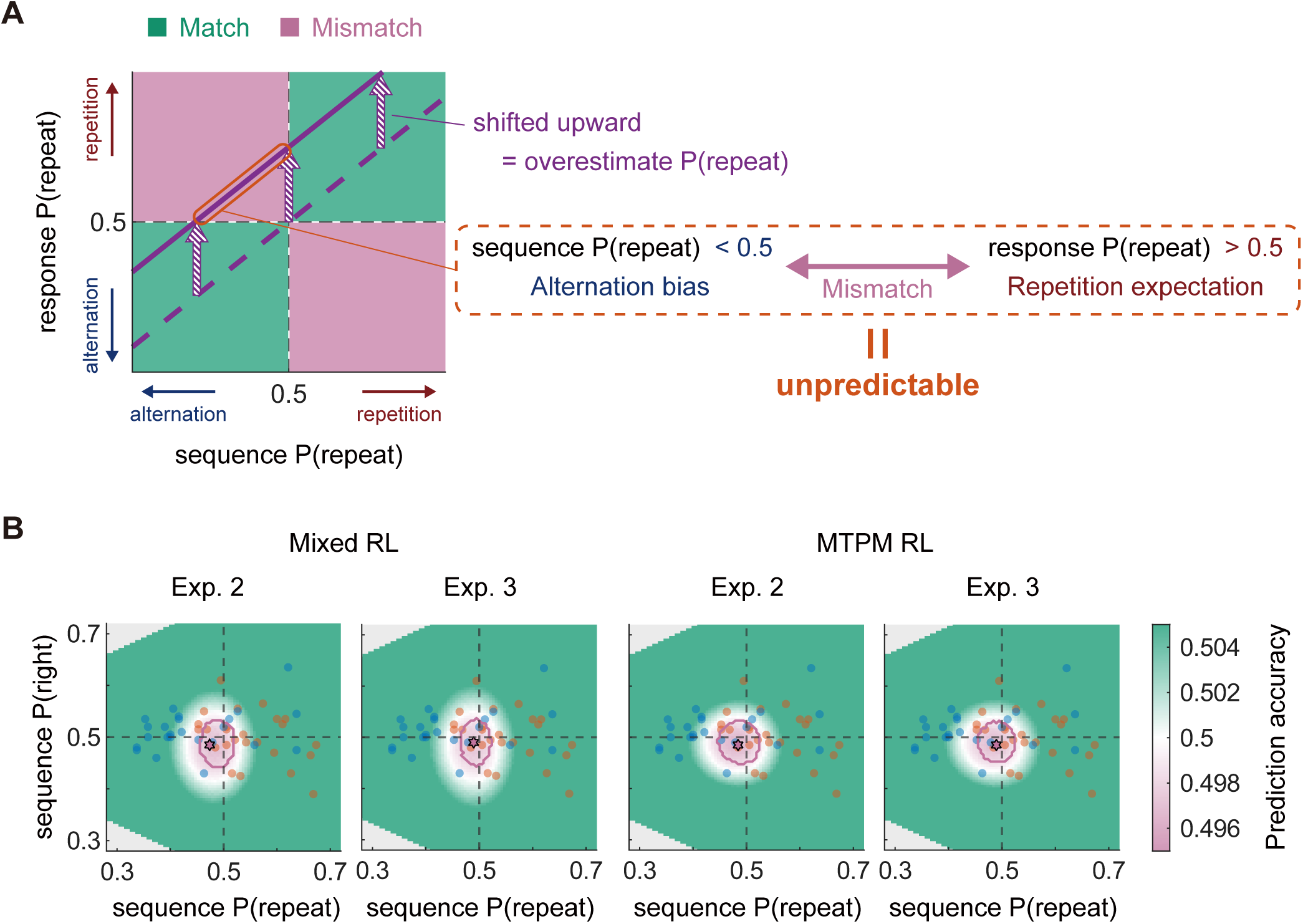
Landscape of human predictability revealing a blind spot shifted toward alternation. (A) Conceptual schematic of how biased calibration shifts the point of minimal predictability. The x-axis shows sequence P(repeat), defined as the probability that the target sequence repeated the immediately preceding item. The y-axis presents response P(repeat), defined as the probability that a human observer predicted such a repetition. If prediction lacked an overall repetition bias (dashed purple line), the prediction line demonstrating the relationship between sequence P(repeat) and response P(repeat) would pass through the neutral point (0.5, 0.5), ensuring that observers correctly expect alternation for alternation-biased sequences and repetition for repetition-biased sequences (matched prediction; green areas). However, as the actual response P(repeat) shifts upward (solid purple line), this line crosses into a mismatch region (pink area) where sequence P(repeat) is below 0.5 but response P(repeat) remains above 0.5. In this mismatch zone, the sequence is biased toward alternation whereas the observer expects repetition, potentially reducing prediction accuracy below the chance level. (B) Model-derived predictability landscapes. Prediction models fitted to the observed data were used to simulate prediction accuracy for finite-length sequences spanning different combinations of sequence P(repeat) and sequence P(right). The columns depict landscapes based on the Mixed RL and MTPM RL models fitted to Experiment 2 or 3. Color indicates simulated prediction accuracy. The pink contour encloses count combinations for which simulated prediction accuracy was significantly below 0.5. The dashed horizontal and vertical lines denote sequence P(right) = 0.5 and sequence P(repeat) = 0.5. Overlaid points show the empirical sequences generated in Experiment 1. Stars indicate the minimum of each simulated accuracy landscape.

We estimated model-derived predictability landscapes to test this possibility. We used two fitted prediction models to capture the observed sequence-level response patterns (Fig. S6A, B): Mixed RL, which learns marginal item probabilities and repetition probabilities separately; and MTPM RL, a reinforcement learning adaptation of the minimal transition probability model^18^, which represents predictions through item-specific transition probabilities (see the Methods). We then simulated how these models predicted finite-length binary sequences spanning different combinations of the sequence P(repeat) and the sequence P(right). Across models and experiments, the predicted accuracy landscape consistently revealed a small region of below-chance performance near, but not exactly at, objective randomness (Fig. 5B). The least predictable points shifted slightly toward alternation in the transition structure (minimum predicted accuracy occurring at sequence P(repeat) = 0.474-0.489). Therefore, the model-derived blind spot for human prediction was not the objectively random point but a slightly alternation-biased region counteracting the human tendency to over-predict repetition.

We then examined whether the same displacement was visible in the actual human prediction data, rather than only in the model-simulated sequences. Quadratic fits relating prediction accuracy to sequence P(repeat) suggested that the minimum of the accuracy curve shifted below 0.5. This shift was clear for the original sequences predicted in Experiment 2 (estimated minimum at sequence P(repeat) = 0.466, CI [0.446, 0.487]; *t* (47) = -3.33, *p* = 0.002, *d* = - 0.33, CI [-0.52, -0.13]) and revealed the same direction for the balanced synthetic sequences predicted in Experiment 3 (estimated minimum at sequence P(repeat) = 0.487, CI [0.472, 0.502]; *t* (47) = −1.74, *p* = 0.089, *d* = -0.13, CI [-0.28, 0.02]) (Fig. S6C). By contrast, the related analysis for sequence P(right) did not reveal a comparable displacement from 0.5 (estimated minimum at sequence P(right) = 0.499, CI [0.471, 0.528]; *t* (47) = −0.05, *p* = 0.961, *d* = -0.02, CI [-0.66, 0.63]) (Fig. S6D). These results signify that the shift in the least predictable point was specific to the transition structure and not the marginal side frequency.

Based on the results, we reinterpret the “unpredictability” of the sequences generated in Experiment 1 from the perspective of a human observer. When evaluated against objective randomness, the Competitive instruction moved generated sequences closer to the transition balance than did the Random instruction, but it also shifted many of them beyond the balance point into repetition. The model-derived landscapes provide a different criterion: the simulated predictability was the lowest not at objective randomness but in a slightly alternation-biased region. Consistently, many Competitive sequences (red points in Fig. 5B) fell outside the model-derived blind spot highlighted by the pink contour, because they tended to occupy the repetition-biased side of the sequence space rather than the slightly alternation-biased region of minimal predictability. Therefore, the failure of Competitive sequences to become maximally unpredictable was not merely a failure to approximate objective randomness. It reflected a mismatch between aspects the generators produced when attempting to evade prediction and the region of the sequence space in which the human predictors were least accurate.

## Discussion

When an opponent’s behavior is directly observable, agents can increase their payoff by deviating from equilibrium mixed-strategy play^4,5^ and exploiting systematic patterns in the opponent’s behavior^27,28^. The present study reveals that humans also depart from randomization in the absence of information about the behavior of the person attempting to predict them—that is, when the potentially exploitable behavior by the opponent is unknown. Yet, this non-random generation pattern did not necessarily make behavior less predictable. We propose that the failure of strategic unpredictability reflected miscalibration in predicting how other humans would predict actions.

One plausible interpretation of the repetition bias in sequences generated to evade prediction is that the generators attempted to exploit the expectations of future observers. Humans often underestimate the probability that the same event will recur after it has just occurred, a phenomenon known as the gambler’s fallacy, widely documented in both laboratory tasks^37–39^ and real-world decisions^40–43^. Therefore, the generators may have projected their own expectations of alternation onto future predictors and deliberately selected repetitions to counteract the presumed expectation. This type of strategic behavior—based on inferences about another person’s reasoning—is formalized in level-k theory and cognitive-hierarchy models ^44–49^. However, as the present study did not directly elicit generator beliefs or reasoning steps, this interpretation remains inferential^49,50^. For instance, some participants may have engaged in deeper recursive reasoning, anticipating that observers would predict their repetition strategies. Nevertheless, such deep strategic reasoning is atypical for most participants^44,47,49,51^.

Human predictors adapted their predictions to the statistical structure of the assigned sequences; however, their predictions were consistently calibrated toward repetition relative to the objective repetition probability. Consequently, slightly alternation-biased sequences created a mismatch between the actual sequence structure and the observers’ expectations: the sequence favored alternation, whereas the observers expected repetition. This mismatch yielded a region in which prediction accuracy could approach or fall below chance. Notably, the present design was static: generators could not dynamically adjust the statistical structure of their sequences in response to the predictors’ behavior^20,52^, offering observers a relative advantage compared with fully interactive settings. The slight shift in the predictability minimum toward alternation aligns with evidence on subjective randomness, which demonstrates that sequences with modest over-alternation often appear most random to human observers^24,53^. Several computational accounts also provide a framework for interpreting this biased subjective randomness^18,54^.

Although human observers adapted their predictions to the sequence statistics, the generators failed to counteract the repetition bias in their sequences, which subsequently remained predictable. Several non-mutually exclusive mechanisms may account for this failure. First, in sequence generation, each element is determined by the generator’s choice. Such internally generated outcomes may provide limited opportunities for the type of externally generated prediction errors that drive learning^2,55,56^, because the outcome tends to match the current action intention^57–59^. Second, the dual requirement of anticipating the predictor’s expectations while maintaining sequence statistics that counter those expectations may have imposed substantial demands on working memory^60,61^. Under such conditions, executive function constraints may have shaped statistical learning^62,63^. Third, when inferring another person’s beliefs^19,64^ and estimating sequence statistics^16,65^, people may rely on partially overlapping prefrontal-centered control and inference networks. Therefore, competition between these neural resources may have compromised the estimation or control of sequence statistics.

More broadly, the current findings show that human predictability is not determined solely by normative solutions or objective behavioral statistics. Rather, it depends on the mapping between the sequence produced by a generator and the biased computations through which an observer predicts that sequence. By separating objective randomness from human predictability, this study suggests that theories of strategic social behavior could be strengthened by considering the imperfect internal models that people use to infer how others interpret their actions.

## Methods

### Participants

A total of 550 individuals participated in the study across three experiments.

Experiment 1 included 50 individuals (female: 25, male: 25; mean age ± SD = 30.9 ± 9.54 years, age range: 18–57 years). We determined the sample size a priori using G*Power^66^. Based on an effect size of *d* = 0.76 from a pilot study assessing the difference between the probability of repeating a choice and the chance level, we determined that 25 participants would be required for the Competitive group to achieve 95% power for a one-sample t-test against the chance level (α = 0.05). Therefore, we recruited 25 participants for the Competitive group and an equal-sized sample of 25 participants for the Random group.

Experiment 2 involved 250 new individuals (female: 145, male: 101, non-binary: 4; mean age ± SD = 27.1 ± 8.82 years, age range: 18–61 years). We determined the sample size a priori using G*Power^66^. Based on an effect size of *d* = 0.40 from a pilot study examining the difference between prediction accuracy and the chance level, 84 participants were required for the Competitive group to achieve 95% power for a one-sample t-test against the chance level (α = 0.05). We decided to recruit 125 participants (five participants for each of the 25 sequences) for the Competitive sequences to ensure that an equal number of participants were assigned to each sequence and account for the variability in statistical features across sequences. An equal-sized sample of 125 participants was recruited for the Random sequences. None of the participants were involved in Experiment 1.

Experiment 3 included 250 new individuals (female: 125, male: 122, non-binary: 3; mean age ± SD = 26.9 ± 9.84 years, age range: 18–68 years). As the prediction task and transition statistics of the target sequences were nearly identical to those in Experiment 2—with the sole exception that the marginal occurrence frequencies of the items were algorithmically balanced—we adopted an identical sample size and allocation strategy. Accordingly, we recruited 125 participants each for the Competitive-like and Random-like sequences, again assigning five participants to each of the 25 sequences per condition. None of the participants were involved in Experiment 1 or 2.

All participants had normal or corrected-to-normal vision, according to self-reports. This study was approved by the Ethical Review Committee for Experimental Research Involving Human Subjects at the Graduate School of Arts and Sciences, the University of Tokyo. All the participants provided informed consent via an online form before study involvement.

### Apparatus

All experiments were conducted using the online experimental platform Gorilla Experiment Builder (https://gorilla.sc). The temporal accuracy of the tool has been validated in previous studies^67,68^. Participants performed the experiment through a web browser on their personal computers.

### Experiment 1: Sequence generation task

First, participants were provided with instructions on how to perform the task. They then completed a 10-trial practice session to become familiar with the procedure. For the main session, participants in the Competitive group were instructed to “select left or right to make the generated sequence unpredictable to an opponent”; meanwhile, those in the Random group were instructed to “select left or right to make the generated sequence as random as possible.” During the main task, participants generated sequences of binary choices (left or right) across 10 blocks of 20 trials each (200 trials in total). In each trial, a cue was presented, and participants had 1500 ms to register their choice via a key press. If no response was detected within the time limit, a warning was displayed and the trial was repeated. Following a valid response, the chosen option was visually highlighted on the screen for 1500 ms. The feedback timing was fixed relative to the trial onset, irrespective of the reaction time. Participants rated their task engagement (the extent to which they had kept their specific instructions in mind) on a 10-point scale at the end of each 20-trial block.

### Stimulus generation for Experiments 2 and 3

In Experiment 2, the exact 50 sequences (200 trials each) generated by the human participants in Experiment 1 (25 Competitive, 25 Random) were used as target sequences.

In Experiment 3, we generated a new set of 50 synthetic “balanced” sequences to isolate the role of transition statistics from marginal side frequencies. Each synthetic sequence was computationally yoked to an original sequence from Experiment 1, for which we employed a large-scale Monte Carlo optimization approach. For each original sequence, we generated extensive candidate sequences strictly constrained to have perfectly balanced marginal frequencies (exactly 100 left and 100 right choices) and an exact match to the first-order transition count (i.e., the total number of repetitions) of the original sequence. Among the candidates addressing these constraints, we chose the single sequence that best approximated the original higher-order run-length statistics. We achieved this by minimizing the absolute difference between the synthetic and original run-length-conditioned repetition probabilities, weighted by the empirical occurrence frequency of each run length in the original data. The synthetic sequences derived from Competitive and Random sequences are referred to as Competitive-like and Random-like sequences, respectively.

In both experiments, the trial order within the blocks was fixed to preserve the temporal transition structure. However, the presentation order of the 10 blocks was randomized for each participant to mitigate potential block-order effects.

### Experiments 2 and 3: Sequence prediction task

The procedure for the prediction task was identical across Experiments 2 and 3. Participants completed a 10-trial practice session before commencing the main session. In the main task, the participants were asked to predict the next item in their assigned target sequence (10 blocks of 20 trials). Participants had 1500 ms to make their prediction (left or right). Following their response, both their predicted choice and the actual target item were displayed for 1500 ms. The feedback timing was set relative to the trial onset, irrespective of the reaction time. Participants were not informed about how or by whom the target sequences were generated, to prevent strategic biasing. At the end of each block, the participants rated their task engagement on a 10-point scale.

### Analysis

All statistical analyses were conducted in MATLAB. We coded the binary sequence items and participant responses as 0 for left and 1 for right. For all transition-based analyses, we excluded transitions across block boundaries; therefore, the first trial of each block was treated as having no preceding item.

For the generated or target sequences, we defined sequence P(right) as the proportion of rightward sequence items. We defined sequence P(repeat) as the proportion of within-block transitions in which the current sequence item repeated the immediately preceding item. As each sequence comprised 10 blocks of 20 trials, we computed sequence P(right) over 200 items, whereas sequence P(repeat) was computed over 190 within-block transitions. For prediction data, we defined response P(right) as the proportion of trials in which the participants predicted a rightward item. Response P(repeat) was defined as the proportion of transition-eligible trials in which participants predicted that the next item would match the immediately preceding target item. We defined prediction accuracy as the proportion of trials in which the participant’s prediction matched the current target item.

In Experiments 2 and 3, five participants were assigned to each target sequence. We used participant-level measures for participant-level tests. For sequence-level analyses, we first computed the values separately for each participant, subsequently averaged across the five participants assigned to the same target sequence. Thus, each target sequence contributed one data point to the sequence-level correlation and regression analyses.

In Experiment 1, the sequence P(right) and the sequence P(repeat) were computed for each participant’s generated sequence. The differences between Competitive and Random sequences were tested using two-tailed independent-samples t-tests. Deviations from objective balance, defined as sequence P(right) = 0.5 or sequence P(repeat) = 0.5, were tested using two-tailed one-sample t-tests. We defined absolute side bias as |sequence P(right)-0.5|, and absolute transition bias as |sequence P(repeat)-0.5|. As these absolute-bias measures were bounded and non-negative, between-sequence class differences were tested using Wilcoxon rank-sum tests. For t-tests, the effect sizes are reported as Cohen’s *d*^69^ with 95% confidence intervals^70^. For the Wilcoxon rank-sum tests, the effect sizes are reported as *r*^71^.

We analyzed block-wise sequence P(right), sequence P(repeat), and engagement ratings using 2 × 10 mixed-design analyses of variance, with instruction as a between-participant factor and block as a within-participant factor. Greenhouse–Geisser correction was applied when appropriate. The mean engagement ratings were also compared between instruction conditions using independent-samples t-tests.

In Experiment 2, we compared prediction accuracy against the 0.5 chance level using two-tailed one-sample t-tests. Differences between Competitive and Random sequences in Experiment 2, and between Competitive-like and Random-like sequences in Experiment 3, were tested using two-tailed independent-samples t-tests. To clarify, Competitive-like and Random-like sequences refer to balanced synthetic sequences derived from the original Competitive and Random sequences, respectively.

We computed response P(repeat) for each participant to assess transition-based prediction. Values > 0.5 indicate a tendency to predict repetition of the immediately preceding target item, while values < 0.5 indicate a tendency to predict alternation. Deviations of response P(repeat) from 0.5 were tested using one-sample t-tests, and sequence-class differences were tested using independent-samples t-tests. For block-wise analyses, we computed the same quantities separately for each block.

We related sequence P(repeat) to response P(repeat) at the sequence level to examine whether the participants’ predictions tracked the transition statistics of individual target sequences. Pearson correlation coefficients were computed separately for Competitive and Random sequence classes, and across all sequences combined. We also fitted linear regression models to sequence-level data to estimate the fitted response P(repeat) at the objective transition balance, sequence P(repeat) = 0.5. For a fitted model,

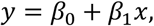

where *x* is sequence P(repeat) and *y* is response P(repeat); the fitted value at *x* = 0.5 was

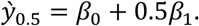

The standard error of y_0.5_ was computed from the coefficient covariance matrix of the fitted regression model. Subsequently, we tested whether y_0.5_ differed from 0.5, using a two-tailed t-test with the residual degrees of freedom of the regression model. Through this analysis, we tested whether response P(repeat) shifted upward relative to sequence P(repeat) at the objective transition balance.

We computed the following formula to directly quantify repetition over-prediction:

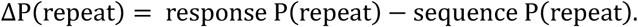

Positive values indicate that participants predicted repetitions more often than repetitions actually occurred in the assigned sequence. We tested these deviations against zero using one-sample t-tests.

The same sequence-level approach was applied to the marginal side frequency. Using Pearson’s correlations and linear regression, we found that sequence P(right) was related to response P(right). These analyses were used to determine whether participants’ predictions were influenced by marginal side frequency information, in addition to transition statistics.

The higher-order transition structure was analyzed using run length-conditioned repetition probabilities. We defined run length as the number of consecutive identical target items immediately preceding the current prediction. Analyses were conducted for run lengths 1–4. For each run length *k*, the run length-conditioned sequence P(repeat) was defined as *P*(*x_t_* = *x_t_*_−1_|*L_t_* = *k*), where *x_t_* is the current target item, *x_t_*_−1_ is the immediately preceding target item, and *L_t_* is the run length immediately preceding trial *t*. The run length-conditioned response P(repeat) was defined as *P*(*y_t_* = *x_t_*_−1_|*L_t_* = *k*), where *y_t_* is the participant’s prediction. Thus, the run length-conditioned sequence P(repeat) describes the empirical higher-order transition structure of the assigned sequence, whereas the run length-conditioned response P(repeat) describes the participant’s expectation that the same item would reoccur after a given run length.

For each run length, we computed the sequence-level correlations between the run length-conditioned sequence P(repeat) and response P(repeat). We then performed linear regressions to estimate the fitted response P(repeat) at sequence P(repeat) = 0.5, using the same coefficient-covariance method described above. Further, we computed response P(repeat) − sequence P(repeat) separately for each run length and tested these deviations against zero to directly examine over-prediction.

In Experiment 3, we used balanced synthetic sequences derived from the original human-generated sequences. For each original sequence, we constructed a synthetic counterpart that balanced the marginal frequencies of the left and right items, while preserving the first-order transition structure and closely matching the run length-conditioned transition statistics of the original sequence. To validate this construction, we compared sequence P(repeat) and the run length-conditioned sequence P(repeat) between each original sequence and its balanced synthetic counterpart. We also compared response P(repeat) between Experiments 2 and 3 at the sequence level to test whether transition-based prediction patterns were preserved when the marginal side-frequency cues were removed.

Further, we fitted two learning models to the participants’ trial-wise predictions and used the fitted parameters to construct model-derived predictability landscapes to estimate where human-like predictions would be least accurate. We focused on two models reproducing the observed sequence-level response patterns for both response P(repeat) and response P(right): a mixed reinforcement learning model (mixed RL) and a minimal transition probability reinforcement learning model (MTPM RL). The mixed RL model represents prediction as a combination of two learned components: a marginal frequency learner, tracking whether the left or right items are more likely overall, and a transition learner, tracking whether the next item tends to repeat or alternate relative to the immediately preceding target item. The MTPM RL model used the item-specific transition probability representation of the minimal transition probability model^18^to estimate the next item conditional on whether the previous target item was left or right. Yet, while the original MTPM was formulated as a Bayesian observer model^18^, we implemented this representation with reinforcement learning updates because the participants in the present task received trial-by-trial feedback regarding the actual target item. Both models included participant-specific response bias and stickiness parameters. The Supplementary Methods detail the full mathematical specifications of the models.

For each participant and model, we estimated the free parameters by minimizing the negative log-likelihood of trial-wise predictions. Model fitting was performed separately for the original sequences predicted in Experiment 2 and balanced synthetic sequences predicted in Experiment 3. We reset the trial history at the beginning of each block, and in transition-dependent predictors, we treated the first trial of each block as transition-ineligible because no immediately preceding target item was available.

First, we validated that the fitted models captured the observed sequence-level response patterns. For each participant, we simulated responses to the empirical target sequences using the fitted parameters and then computed the simulated response P(repeat) and response P(right). These values were averaged across the five participants assigned to each target sequence and compared with the corresponding observed sequence-level values.

Next, we generated finite-length binary target sequences spanning the space of sequence P(right) and sequence P(repeat) to construct the predictability landscapes. Each simulated target sequence comprised 200 trials arranged in 10 blocks of 20 trials, matching the experimental tasks. Sequence P(right) was defined as the number of rightward target items divided by 200. Sequence P(repeat) was defined as the number of repeated within-block transitions divided by 190, as the first trial of each block had no immediately preceding target item. We enumerated the exact finite-count combinations using right-item counts from 0 to 200 and repeated transition counts from 0 to 190. We excluded count combinations that were mathematically impossible for 10 blocks of 20 trials.

For each feasible count combination, we sampled the target sequences exactly matching the specified number of rightward items and repeated transitions. Exact sampling was implemented using a dynamic programming sampler, ensuring that the generated sequences matched the requested finite counts rather than only approximating the nominal probabilities. For the final landscapes, we sampled 1,000 target sequences for each feasible count combination. For each sampled sequence, we simulated trial-wise predictions using the fitted model parameters for each participant. For each count combination and participant, we computed prediction accuracy, response P(right) and response P(repeat). Values were first averaged across sampled sequences within each count combination and then across participants to obtain the model-derived landscape for each model and experiment.

To identify below-chance regions in the landscapes, we tested the simulated prediction accuracy against 0.5 at each feasible count combination using one-tailed one-sample t-tests across fitted participants, assessing whether the simulated accuracy was below the chance level. The point of minimum predicted accuracy was defined as the feasible count combination with the lowest mean simulated prediction accuracy.

We also tested whether a shift in the least predictable point was visible in the empirical sequence-level data. For each target sequence, the prediction accuracy was averaged across five participants assigned to the particular sequence. Next, we fitted ordinary least-squares quadratic regressions relating sequence-level prediction accuracy to either sequence P(repeat) or sequence P(right), distinctly for the original sequences in Experiment 2 and the balanced synthetic sequences in Experiment 3:

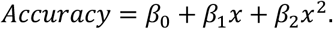

The x-coordinate of the vertex was estimated as follows:

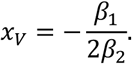

The standard error and 95% confidence interval of *x_V_* were estimated using the delta method, based on the coefficient covariance matrix of the fitted regression model. The gradient of *x_V_* vis-à-vis (*β*_0_, *β*_1_, *β*_2_) was

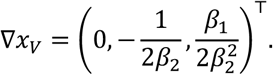

Thus,

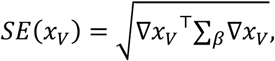

where ∑*_β_* is the coefficient covariance matrix. We tested whether the estimated minimum differed from the objective balance using a two-tailed t-test of

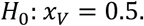

The same procedure was applied to sequence P(repeat) and sequence P(right). Through this analysis, we tested whether the empirical prediction accuracy was minimized at the objective transition balance or at a shifted point in the sequence statistic space. Effect sizes for the vertex tests were computed by standardizing x_V-0.5 by the standard deviation of the observed sequence-level predictor.

For all analyses involving multiple comparisons across blocks (1–10), run lengths (1–4), and the grid of feasible count combinations in the predictability landscapes, we adjusted p-values using the Benjamini–Hochberg false discovery rate procedure to control the false discovery rate^72^. The significance threshold was set at α = 0.05.

## Author contributions

K.Y. and K.T. conceptualized the study. K.Y. and K.T. developed the methodology. K.Y. collected the data and performed formal analyses. K.Y. wrote the original draft. All authors reviewed and edited the manuscript. K.T. and K.K. supervised the project.

## Acknowledgments

We thank the members of the Kudo Laboratory for their helpful discussions and comments, as well as the administrative staff for their support. This work was supported by KAKENHI (JP22K17673, JP24H01434, JP25H01237, and JP25KJ1145) from the Japan Society for the Promotion of Science (JSPS), Strategic Basic Research Programs ACT-X (JPMJAX24LA) from the Japan Science and Technology Agency, and the Kubota endowed fund from Kubota Corporation.

## Data availability

The data and code supporting the findings of this study are available from the authors upon reasonable request.

---

**Figure S1.**
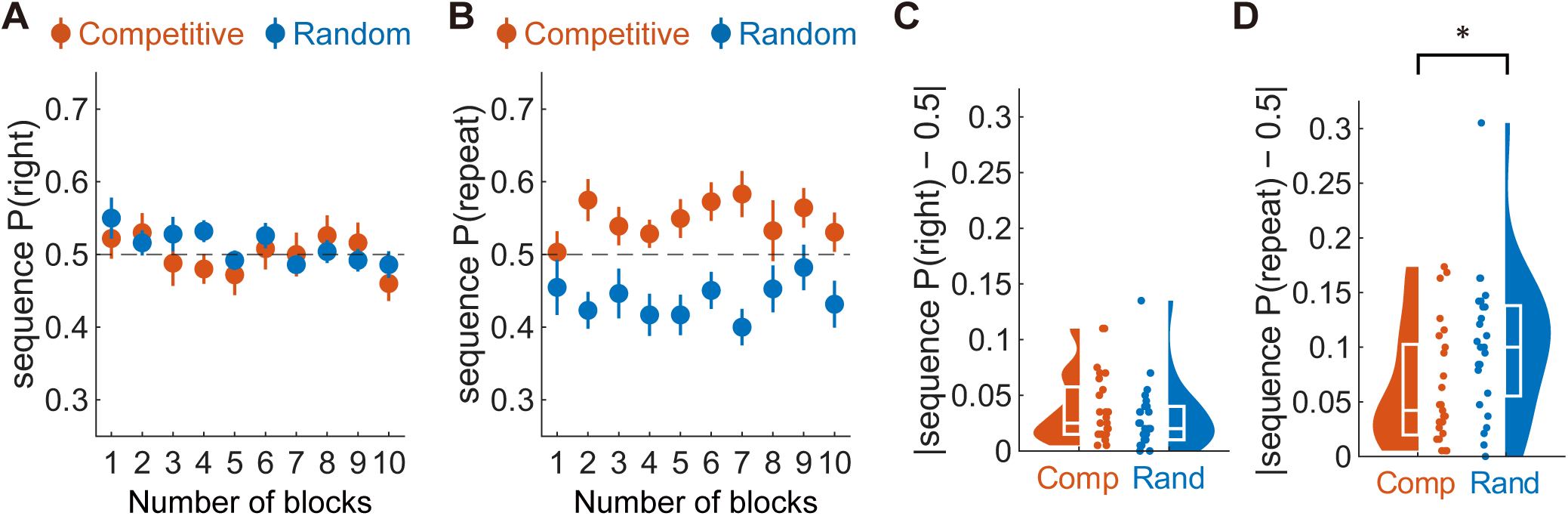
Marginal side use and block-wise dynamics not accounting for the transition bias in Experiment 1. (A) Block-wise sequence P(right) for Competitive and Random sequences. Sequence P(right) denotes the probability that an item in the generated sequence was rightward. Points show group means separately for each block; error bars indicate ±1 SEM. The horizontal dashed line denotes balanced side use, sequence P(right) = 0.5. (B) Block-wise sequence P(repeat) for Competitive and Random sequences. Sequence P(repeat) denotes the probability that a sequence item repeated the immediately preceding item. Points show group means separately for each block; error bars indicate ±1 SEM. The horizontal dashed line signifies the chance-level transition balance, sequence P(repeat) = 0.5. (C) Overall magnitude of side bias, defined as |sequence P(right) − 0.5|, collapsed across blocks. Larger values indicate stronger preference for one side, irrespective of direction. (D) Overall magnitude of transition bias, defined as |sequence P(repeat) − 0.5|, collapsed across blocks. Larger values denote stronger deviation from first-order transition balance, irrespective of whether the deviation reflects repetition or alternation. In Panels C and D, half-violin plots show participant-level distributions, embedded box plots show the median and interquartile range, and the dots indicate individual participants. Black asterisks denote between-sequence class differences. \**p* < 0.05.

**Figure S2.**
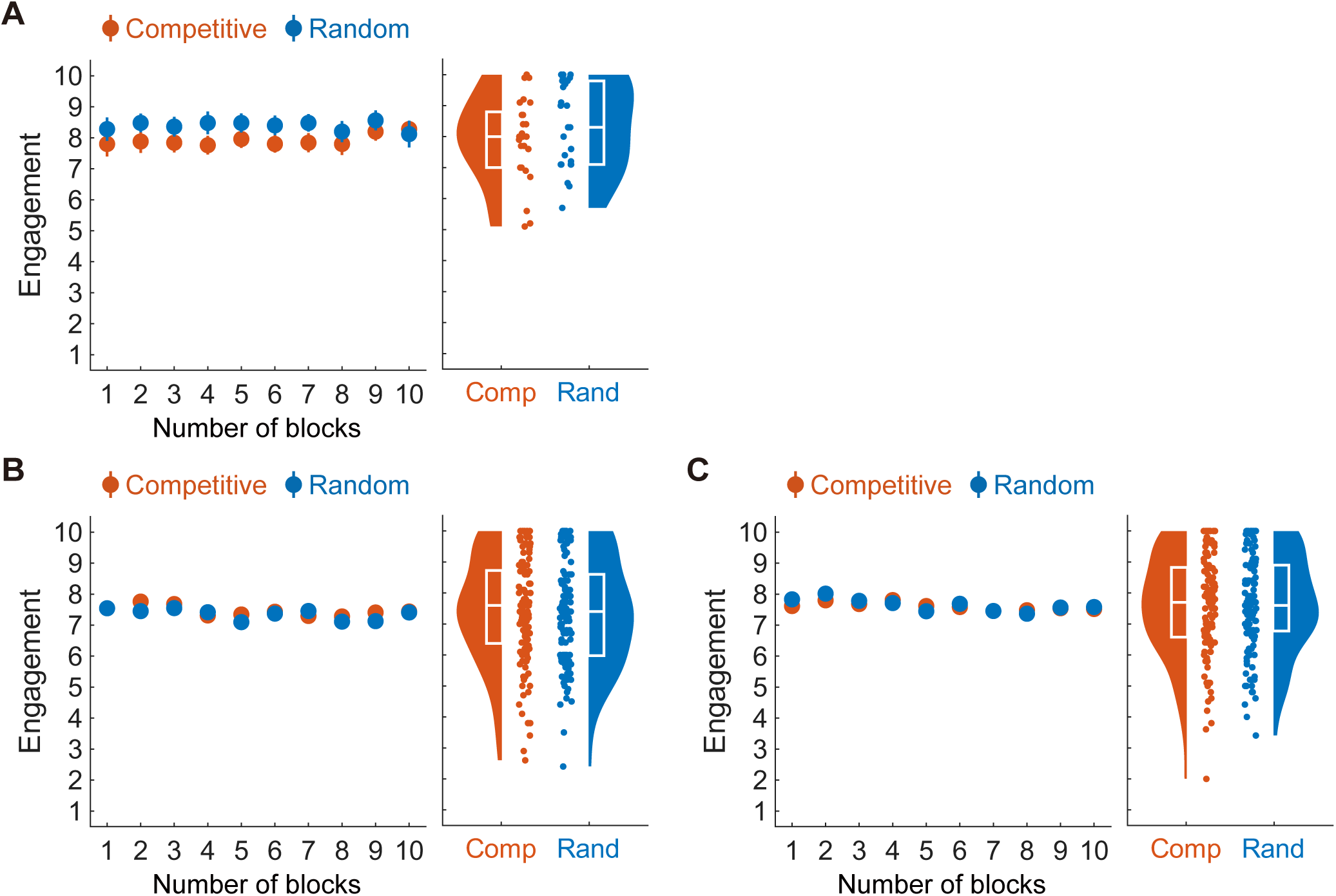
Comparable task engagement between groups across experiments. (A) Self-reported task engagement in Experiment 1, with participants generating sequences based on either a Competitive or Random instruction. (B) Self-reported task engagement in Experiment 2, with participants predicting the original Competitive or Random sequence generated in Experiment 1. (C) Self-reported task engagement in Experiment 3, with participants predicting the balanced Competitive-like or Random-like sequences. In each panel, the left plot shows engagement ratings across the 10 blocks; points indicate group means and error bars indicate ±1 SEM. The right plot shows the participant-level mean engagement ratings collapsed across blocks, half-violin plots show the distribution, embedded box plots show the median and interquartile range, and the dots indicate individual participants.

**Figure S3.**
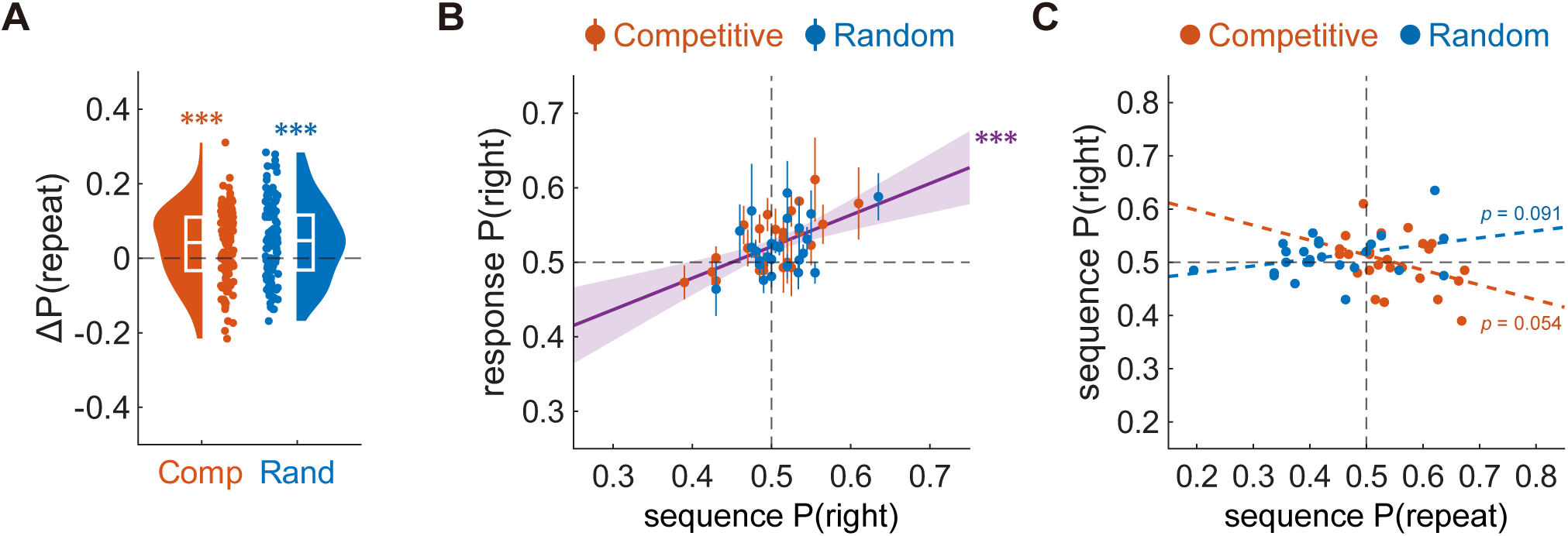
Sequence-level transition and marginal tracking in original human-generated sequences in Experiment 2. (A) Repetition over-prediction relative to the assigned sequence. Values show response P(repeat) − sequence P(repeat), collapsed across blocks. Positive values show that participants predicted repetitions more often than repetitions actually occurred in the assigned sequence. (B) Relationship between sequence P(right) and response P(right). Sequence P(right) denotes the probability that the target item was rightward; response P(right) denotes the probability that participants predicted a rightward item. Each point represents one target sequence; error bars indicate ±1 SEM across the five participants assigned to the particular sequence. The purple line shows the linear fit across all sequences; shading indicates the 95% confidence interval. Horizontal and vertical dashed lines indicate P(right) = 0.5. (C) Relationship between sequence P(repeat) and sequence P(right). Each point represents one target sequence. Dashed lines show separate linear fits for Competitive and Random sequences. Red and blue indicate Competitive and Random sequences, respectively. Asterisks denote one-sample differences from zero in panel A. \*\*\**p* < 0.001.

**Figure S4.**
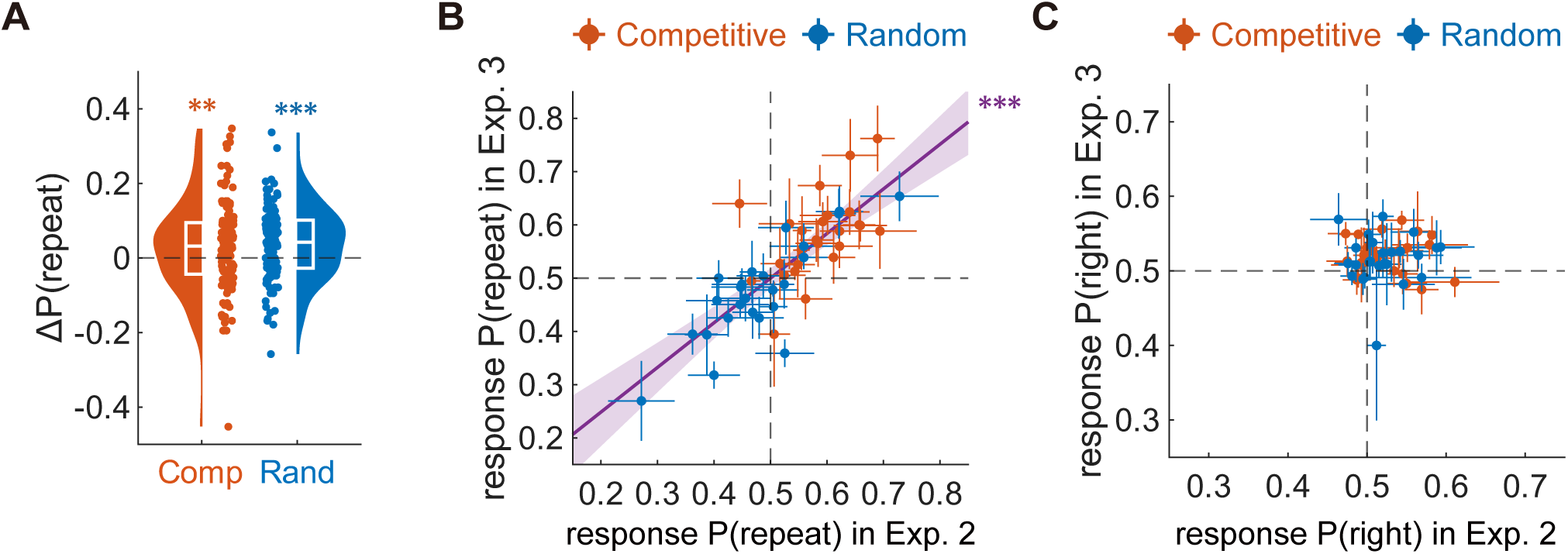
Sequence-level tracking preserved for transition but not marginal statistics in synthetic sequences. (A) Repetition over-prediction relative to the assigned balanced sequence. Values show response P(repeat) − sequence P(repeat), collapsed across blocks. Positive values show that participants predicted repetitions more often than repetitions actually occurred in the assigned sequence. (B) Sequence-level response P(repeat) is preserved across Experiments 2 and 3. Each point represents one sequence pair: an original sequence predicted in Experiment 2 and its balanced synthetic counterpart predicted in Experiment 3. The x-axis shows response P(repeat) in Experiment 2; the y-axis shows response P(repeat) in Experiment 3. Error bars indicate ±1 SEM across the five participants assigned to each sequence in each experiment. (C) Sequence-level response P(right) is not preserved across Experiments 2 and 3. Plotting conventions are the same as in Panel B, except that the axes show response P(right). Red and blue indicate the Competitive-like and Random-like sequences, respectively. Asterisks denote one-sample differences from zero in Panel A. \*\**p* < 0.01, \*\*\**p* < 0.001.

**Figure S5.**
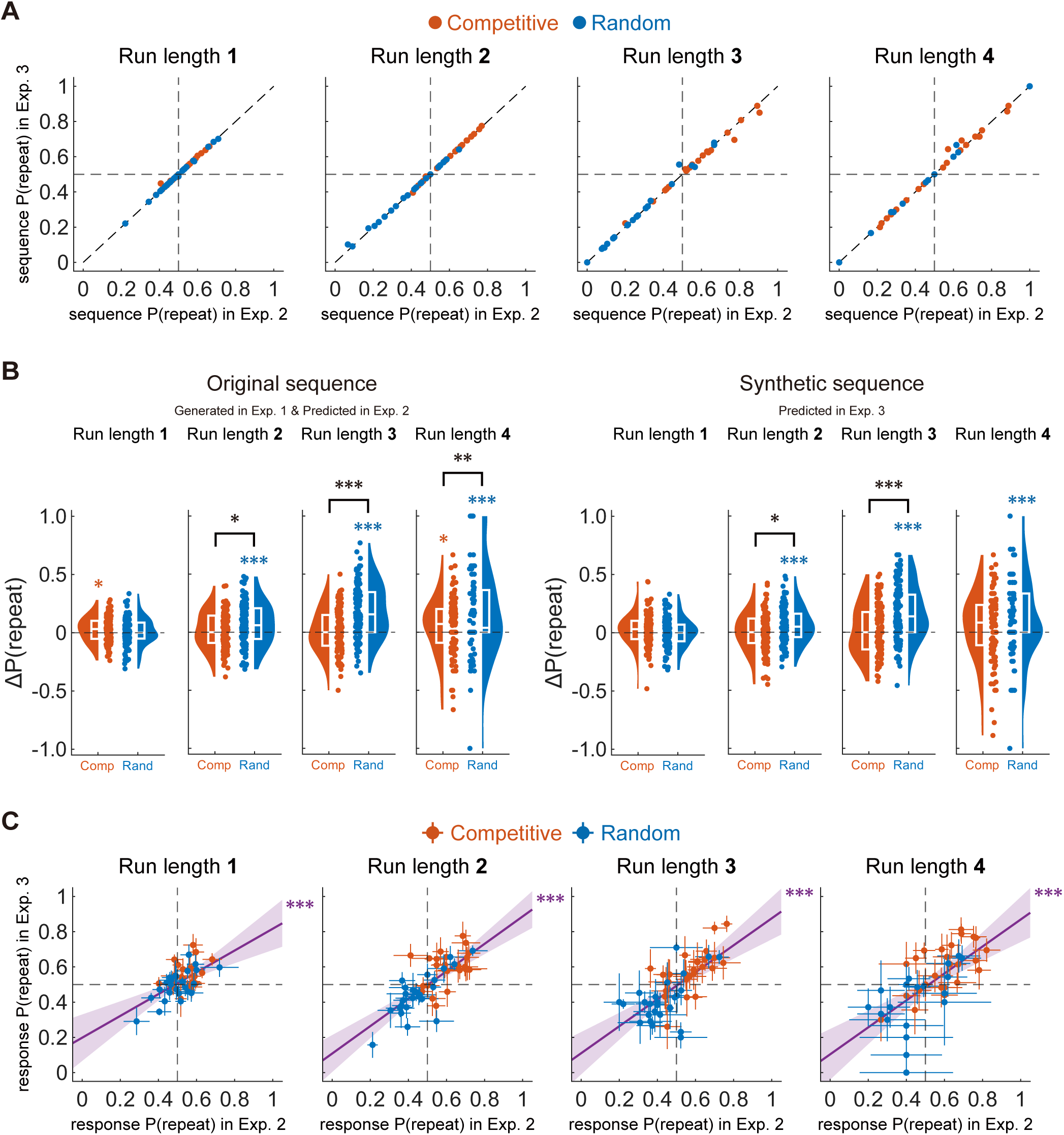
Validation of run-length-conditioned transition statistics and robustness of higher-order predictions. (A) Preservation of run-length-conditioned transition statistics in the synthetic sequences. Each panel depicts the relationship between sequence P(repeat) in the original sequences predicted in Experiment 2 and sequence P(repeat) in the corresponding balanced synthetic sequences predicted in Experiment 3, separately for each run length. Each point represents one sequence pair. The diagonal dashed line denotes the identity line; horizontal and vertical dashed lines indicate P(repeat) = 0.5. (B) Run length-conditioned repetition over-prediction relative to the assigned sequence. Values show response P(repeat) − sequence P(repeat), separately for original sequences in Experiment 2 and balanced synthetic sequences in Experiment 3. Positive values demonstrate that participants predicted repetitions more often than repetitions actually occurred after the particular run length in the assigned sequence. Half-violin plots show participant-level distributions; embedded box plots present the median and interquartile range; dots indicate individual participants. (C) Run length-conditioned response P(repeat) is preserved across Experiments 2 and 3. Each panel presents the relationship between response P(repeat) for the original sequences predicted in Experiment 2 and response P(repeat) for the corresponding balanced synthetic sequences predicted in Experiment 3, separately for each run length. Each point represents one sequence pair; error bars indicate ±1 SEM across the five participants assigned to each sequence in each experiment. The purple line indicates the linear fit across all sequences; shading captures the 95% confidence interval. Horizontal and vertical dashed lines indicate P(repeat) = 0.5. Red and blue indicate Competitive/Competitive-like and Random/Random-like sequences, respectively. Black brackets denote between-sequence class differences, and red and blue asterisks denote one-sample differences from zero within Competitive/Competitive-like and Random/Random-like sequences, respectively. \**p* < 0.05, \*\**p* < 0.01, \*\*\**p* < 0.001.

**Figure S6.**
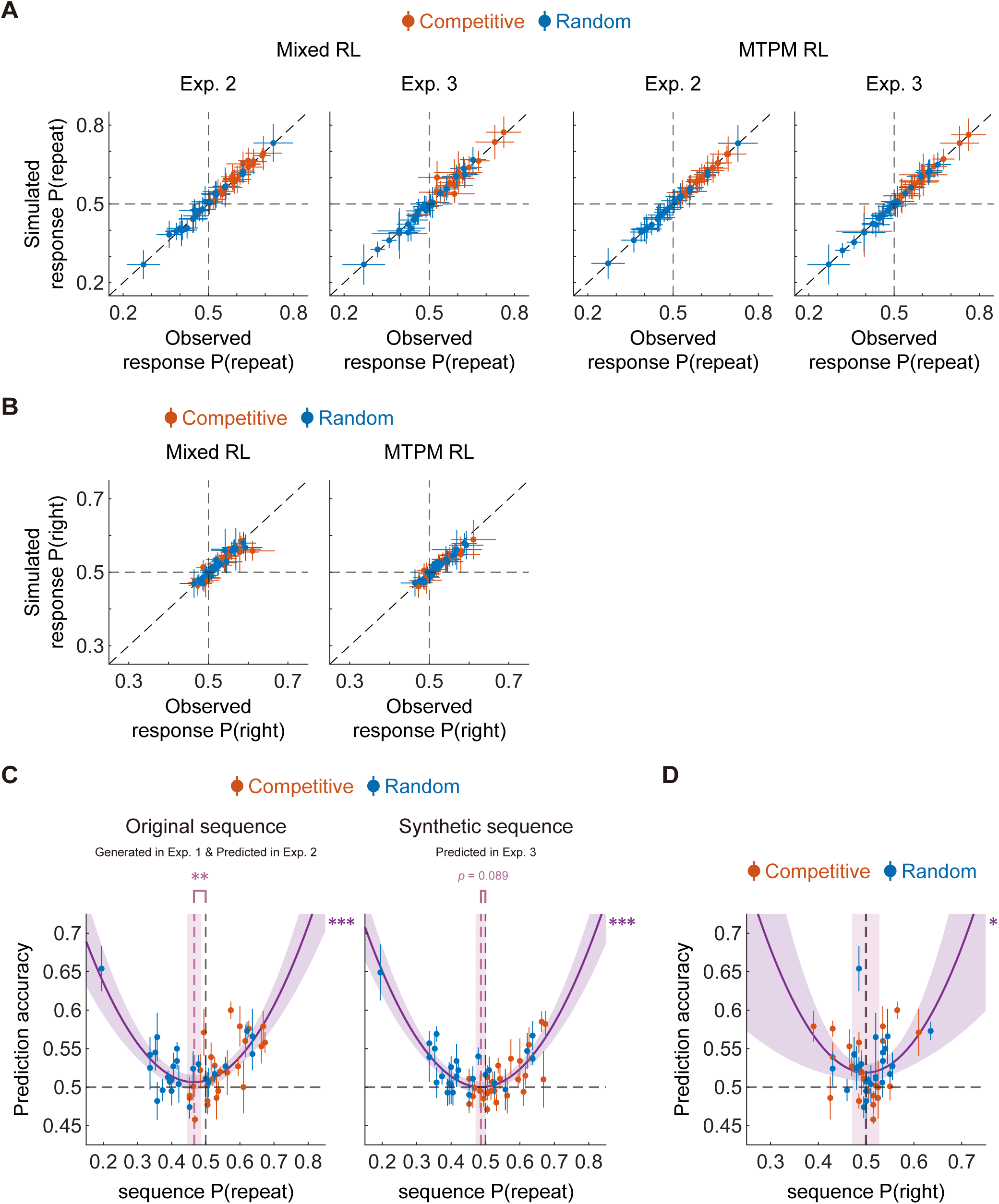
Computational model validation and empirical shifts in minimal predictability. (A) Model validation for response P(repeat). Observed sequence-level response P(repeat) is plotted against simulated response P(repeat) for the Mixed RL and MTPM RL models, separately for the original sequences predicted in Experiment 2 and the balanced synthetic sequences predicted in Experiment 3. Each point represents one sequence; error bars indicate ±1 SEM across participants assigned to the particular sequence. The diagonal dashed line denotes the identity line. (B) Model validation for response P(right). Plotting conventions are the same as in Panel A, except that the axes show observed and simulated response P(right). (C) Empirical relationship between sequence P(repeat) and prediction accuracy. Quadratic fits were used to estimate the sequence P(repeat) value at which prediction accuracy was minimized, separately for the original sequences in Experiment 2 and the balanced synthetic sequences in Experiment 3. The vertical pink dashed line indicates the estimated minimum; the shaded pink band denotes its 95% confidence interval. The black vertical dashed line shows sequence P(repeat) = 0.5, and the horizontal dashed line indicates chance-level accuracy. (D) Empirical relationship between sequence P(right) and prediction accuracy. Plotting conventions are the same as in Panel C. Red and blue indicate Competitive/Competitive-like and Random/Random-like sequences, respectively. Purple asterisks indicate the significance of the quadratic association between the sequence statistic and prediction accuracy. \**p* < 0.05, \*\**p* < 0.01, \*\*\**p* < 0.001.

