## Supplementary Methods for "Trying to be unpredictable even against unobserved opponents shifts humans away from randomness but not from human predictability"

For both models, the target item on trial *t* was denoted by $x_t \in \{0, 1\}$, where 0 indicates left and 1 indicates right. The participant's prediction was denoted by $y_t \in \{0, 1\}$, where 1 indicates a rightward prediction. The logistic function was defined as follows:

$$\sigma(z) = \frac{1}{1 + \exp(-z)}.$$

For any binary variable $u \in \{0, 1\}$, signed coding was defined as follows:

$$\tilde{u} = 2u - 1.$$

We use $\mathbb{I}(\cdot)$ to denote an indicator function, equaling 1 when the statement in parentheses is true and 0 otherwise. A repetition prediction was defined by the binary variable

$$A_t = \mathbb{I}(y_t = x_{t-1}),$$

which equals 1 when the participant predicted the same item as the immediately preceding target item and 0 when the participant predicted the opposite item. To index repetition and alternation values, we also defined the corresponding categorical prediction label $C_t$, where $C_t = \text{rep}$ if $A_t = 1$ and $C_t = \text{alt}$ if $A_t = 0$. The actual transition of the target sequence was defined as follows:

$$T_t = \mathbb{I}(x_t = x_{t-1}),$$

which equals 1 for a repetition transition and 0 for an alternation transition. Transition-dependent quantities were only defined for trials with immediately preceding target items in the same block. When a signed repetition-prediction history term was required, we used

$$\widetilde{A_t} = 2A_t - 1.$$

We also defined a signed side-history term based on the previous target item,

$$\widetilde{x_{t-1}} = 2x_{t-1} - 1,$$

and a motor-level repetition history term,

$$M_t = \mathbb{I}(y_t = x_t),$$

with signed form

$$\widetilde{M_t} = 2M_t - 1.$$

Here, $M_t = 1$ indicates that that the participant's prediction in trial *t* matched the target item in the particular trial; consistently, in the next trial, it coded whether the participant's previous response was aligned with the previous target item. History-dependent signed terms were set to zero when a relevant previous response or target item was unavailable.

The Mixed RL model combined a marginal frequency learner and transition learner. The marginal frequency learner estimated the predictive value of choosing the left or right option. Let $Q_L(t)$ and $Q_R(t)$ denote the learned values of predicting left and right at trial *t*. The marginal frequency component generated the probability of predicting the right option as:

$$P_t^{freq}(R) = \sigma\big(\beta_{freq}\big(Q_R(t) - Q_L(t)\big) + b_{LR} + s_{LR}\tilde{y}_{t-1}\big),$$

where $\beta_{freq}$ is the inverse temperature parameter for marginal side values, $b_{LR}$ is a side bias parameter, and $s_{LR}$ captures response stickiness. The stickiness terms were set to 0 when the relevant previous response was unavailable. The chosen marginal value was updated according to a Rescorla–Wagner rule:

$$Q_{y_t}(t+1) = Q_{y_t}(t) + \alpha_{freq}\left(\mathbb{I}(y_t = x_t) - Q_{y_t}(t)\right),$$

$$Q_{y_t} = \begin{cases} Q_L \ (y_t = 0) \\ Q_R \ (y_t = 1) \end{cases},$$

where $\alpha_{freq}$ is the learning rate for the marginal-frequency component. The unchosen value remained unchanged.

The transition learner estimated the predictive value of repetition and alternation. Let $Q_{rep}(t)$ and $Q_{alt}(t)$ denote the learned values for predicting repetition and alternation, respectively. The probability of predicting repetition was

$$P_t^{rep} = \sigma\left(\beta_{tr}\left(Q_{rep}(t) - Q_{alt}(t)\right) + b_{rep} + s_{rep}\tilde{A}_{t-1} + b_{LR}\tilde{x}_{t-1} + s_{LR}\tilde{M}_{t-1}\right),$$

where $\beta_{tr}$ is the inverse temperature parameter for the transition values, $b_{rep}$ is a repetition bias parameter, and $s_{rep}$ captures the repetition choice stickiness. The transition component was converted into the probability of predicting the right option, as follows:

$$P_t^{trans}(R) = x_{t-1}P_t^{rep} + (1 - x_{t-1})(1 - P_t^{rep}).$$

Thus, if the previous target item was rightward, a repetition prediction corresponded to predicting the right option; if the previous target item was leftward, an alternation prediction corresponded to predicting the right option. The transition values were updated according to the following formula:

$$Q_{C_t}(t+1) = Q_{C_t}(t) + \alpha_{tr}\left(\mathbb{I}(A_t = T_t) - Q_{C_t}(t)\right).$$

The unchosen value remained unchanged. The same transition learning rate was used for repetition and alternation in the Mixed RL model used for predictability landscape analysis.

The final probability of predicting right was a weighted mixture of the marginal frequency and transition components:

$$P_t(R) = wP_t^{freq}(R) + (1 - w)P_t^{trans}(R),$$

$$w = \sigma(\omega).$$

The parameter w controls the relative contribution of the marginal frequency learner. In the transition ineligible trials, the transition component was set to chance.

The MTPM RL model learned item-specific first-order transition probabilities. Specifically, it estimated the probability of a right item following a left item and the probability of a left item following a right item:

$$q_{1|0}(t) \approx P(x_t = 1 | x_{t-1} = 0),$$

$$q_{0|1}(t) \approx P(x_t = 0 | x_{t-1} = 1).$$

Both estimates were initialized at 0.5. Given the previous target item, in the model, the belief that the next item would repeat was captured by:

$$\hat{p}_t^{rep} = \begin{cases} 1 - q_{1|0}(t) \ (x_{t-1} = 0) \\ 1 - q_{0|1}(t) \ (x_{t-1} = 1) \end{cases}.$$

The probability of predicting repetition was then

$$P_t^{rep} = \sigma\left(\beta_{TP}\left(\hat{p}_t^{rep} - 0.5\right) + b_{rep} + s_{rep}\tilde{a}_{t-1} + b_{LR}\tilde{x}_{t-1} + s_{LR}\tilde{m}_{t-1}\right),$$

where $\beta_{TP}$ maps the learned transition-probability estimate onto the response probability. This repetition prediction probability was converted into a probability of predicting right as

$$P_t(R) = x_{t-1}P_t^{rep} + (1 - x_{t-1})(1 - P_t^{rep}).$$

After observing the target item, the relevant transition estimate was updated using a Rescorla–Wagner rule. If the previous target item was leftward,

$$q_{1|0}(t+1) = q_{1|0}(t) + \alpha_{TP}\left(x_t - q_{1|0}(t)\right).$$

If the previous target item was rightward,

$$q_{0|1}(t+1) = q_{0|1}(t) + \alpha_{TP}\left((1 - x_t) - q_{0|1}(t)\right).$$

The other transition estimate remained unchanged in the particular trial. Here, $\alpha_{TP}$ is the learning rate for item-specific transition probabilities.

For each participant and model, the free parameters were estimated by minimizing the negative log-likelihood of trial-wise predictions. The likelihood of a response in trial *t* was

$$P(y_t|\theta) = P_t(R)^{y_t}(1 - P_t(R))^{1-y_t},$$

where $\theta$ denotes the model parameters. The negative log-likelihood was

$$\mathcal{L}(\theta) = -\sum_{t\in\mathcal{T}}(y_t \log P_t(R) + (1 - y_t)\log(1 - P_t(R))),$$

where $\mathcal{T}$ denotes the likelihood-eligible trials. The trial history was reset at the beginning of each block. Transition-dependent predictors required an immediately preceding target item and were, therefore, ineligible for the first trial of each block.

For posterior predictive checks and predictability landscape simulations, the fitted parameters were used to generate simulated responses according to

$$y_t^{sim} \sim \mathrm{Bernoulli}\left(P_t(R)\right).$$

Model states were updated sequentially using the same update rules followed in model fitting.
